# Widespread exon definition is promoted by the U1 snRNP 70K subunit and requires sites of contact with RNA Polymerase II

**DOI:** 10.64898/2026.09.07.749963

**Authors:** Nova Fong, Ryan Sheridan, Benjamin Erickson, Rui Zhao, Aaron A. Hoskins, J. Matthew Taliaferro, David L. Bentley

## Abstract

Much pre-mRNA splicing initiates co-transcriptionally, but its mechanism is unclear. We investigated this process by degron depletion of the U1-70K subunit of U1 snRNP that contacts transcribing RNA polymerase II (RNAPII). U1-70K deficiency caused extensive exon skipping consistent with widespread exon definition. Surprisingly, exons with the strongest 5’ splice sites (SSs) are skipped. Furthermore, strengthening of non-canonical 5’ splice sites with a complementary mutant U1 snRNA or a splice modifying drug induced exon skipping in response to U1-70K depletion. We suggest that U1-70K plays a previously unappreciated role in destabilizing the U1-5’SS interaction to facilitate the transition to the U6-5’SS interaction required for productive splicing following exon definition. Consistent with this idea, knock-down of Prp28/DDX23 that exchanges U1 for U6 at the 5’SS also promotes exon skipping mimicking U1-70K depletion. Replacing WT U1-70K in degron-containing cells with a mutant in the RNAPII interface caused widespread exon skipping. We propose that frequent exon definition occurs at the transcription elongation complex and is enabled by U1-70K contact with transcribing RNAPII.

## Introduction

pre-mRNA splicing frequently occurs on elongating nascent transcripts (Carrillo Oesterreich et al. 2010; Khodor et al. 2011; Bentley 2014) but how it is coordinated with ongoing transcription and how it may differ from splicing uncoupled from transcription are not well understood (Shenasa and Bentley 2023; Carrocci and Neugebauer 2024). The potential significance of co-transcriptional splice site recognition is underscored by two recently characterized interactions between RNAPII and a) U1 snRNP (Zhang et al. 2021) that recognizes the 5’ splice site (5’SS), and b) U2AF that recognizes the 3’SS (Shao et al. 2025). U1 snRNP makes contacts with transcribing RNAPII in a way that is compatible with recognition of the 5’SS in the nascent transcript. Specifically, the RRM domain of the U1-70K (SNRNP70) subunit of U1 snRNP contacts the Rpb2 and Rpb12 subunits of the polymerase near the RNA exit channel (Zhang et al. 2021). This discovery begs the questions of how contact between U1 snRNP and RNAPII affects transcription and co-transcriptional assembly of splicing complexes.

*In vitro*, splicing of pre-mRNAs can occur by intron definition where 5’ and 3’ SSs pair across the intron, or by exon definition where initial cross exon pairing is subsequently converted to cross-intron pairing (Schneider et al. 2010; Boesler et al. 2015; Li et al. 2019; Zhang et al. 2024b). One attractive idea is that U1 snRNP-5’SS complexes on the RNAPII surface are positioned to make the cross-exon contacts with U2AF-3’SS complexes that are co-localized on the polymerase via binding of U2AF1 to Rpb9 (Shao et al. 2025). In this way the RNA polymerase could serve as a hub to facilitate co-transcriptional exon definition (Shenasa and Bentley 2023). However, it remains to be determined whether exon definition is a widely used mechanism of SS recognition *in vivo*. The *in vivo* significance of exon definition has been called into question by the discovery of superfast splicing that occurs well before the 5’SS of the downstream exon has been transcribed, thereby precluding exon definition (Oesterreich et al. 2016; Zeng et al. 2022). In contrast, other studies suggest that much co-transcriptional splicing is completed after RNAPII has extended well beyond the 3’ exon when exon definition is possible (Drexler et al. 2020; Gildea et al. 2022). The *in vivo* relevance of the exon definition complexes characterized *in vitro* (Schneider et al. 2010), has also been questioned. A recently proposed alternative model is that instead of independent recognition of upstream and downstream 5’SS’s by two U1snRNPs followed by conversion of cross-exon to cross-intron complexes, a single U1 snRNP is transferred from the upstream 5’ SS to the downstream 5’SS by a "relay" that is coupled to activation of the spliceosome on the upstream intron (Yoon et al. 2026). Notably, the U1 relay model predicts that a strong downstream 5’SS will favor transfer of the U1 snRNP and inclusion of the adjacent exon. Exon definition can be distinguished from intron definition by mutating the 5’ SS of the downstream exon, which results in intron retention if intron definition operates, or exon skipping if exon definition operates (Robberson et al. 1990; Kuo et al. 1991). The significance of exon definition is supported by the effects of naturally occurring 5’SS mutations (Krawczak et al. 2007), however, the strategy of experimentally mutating 5’SSs to diagnose intron versus exon definition cannot be applied *in vivo* transcriptome-wide. As an alternative, we attempted to perturb 5’SS recognition generally by depleting cells of U1-70K, which is unique to U1 snRNP, and asking how splicing was affected.

Some evidence already exists that U1-70K may promote exon definition. For example, in addition to RNAPII, U1-70K also contacts SRSF1 that facilitates formation of cross-exon complexes (Wu and Maniatis 1993; Kohtz et al. 1994; Hertel and Graveley 2005; Cho et al. 2011; Jobbins et al. 2022; Paul et al. 2024). U1-70K is also critical for formation of exon definition complexes *in vitro* (Rogalska et al. 2023) although its precise role in this process is unknown. Re-constituted U1 snRNPs lacking U1-70K can support splicing but have reduced affinity for the 5’ SS likely because U1-70K contacts U1-C that stabilizes the U1-5’SS interaction (Will et al. 1996; Kondo et al. 2015; Rogalska et al. 2023).

U1-5’SS complexes are disrupted by the DDX23/PRP28 helicase when U6 replaces U1 at the 5’SS as the pre-B complex transitions to the B complex (Staley and Guthrie 1999; Wilkinson et al. 2020). Interestingly, yeast U1 snRNP increases the dissociation rate of U1-5’SS complexes (Hansen et al. 2022) suggesting a possible role for this snRNP in mobilizing the 5’SS for transfer to U6. Like U1-70K, DDX23 also interacts with SRSF1 potentially implicating this helicase in exon definition (Segovia et al. 2024).

How transcription elongation may be influenced by the splicing machinery to optimize co-transcriptional splicing is poorly understood. Notably U1 snRNP stimulates transcription elongation, possibly by contacting RNAPII directly, although the mechanism is unclear. Antisense morpholino oligonucleotide (AMO) disruption of U1 snRNP (Feng et al. 2023) decelerates transcription, and promotes premature termination (Mimoso and Adelman 2023). Transcriptional pausing frequently punctuates elongation and could facilitate recognition of a SS while it is proximal to the polymerase. Transcriptional pauses near 5’ and 3’ SSs have been reported (Alexander et al. 2010; Carrillo Oesterreich et al. 2010; Oesterreich et al. 2011; Nojima et al. 2015; Milligan et al. 2017) but their validity has been questioned because it is difficult to distinguish RNA 3’ ends at pause sites from splicing intermediates (Sheridan et al. 2019; Reimer et al. 2021; Shenasa and Bentley 2023) that co-purify with RNAPII. Whether U1 snRNP affects pausing either generally or at SSs is not known.

Here we depleted U1-70K *in vivo* using an inducible protein degradation system. U1-70K degradation is expected to prevent direct interaction of U1 snRNP with the polymerase and to generally impair 5’SS recognition. This approach allowed us to investigate the importance of U1 snRNP contact with RNAPII for transcription elongation and co-transcriptional splicing. NET-seq revealed that RNAPII pauses preferentially one base downstream of the 5’SS, but this process is independent of U1070K. U1-70K depletion had a major effect on splicing however, manifested by widespread exon skipping with little intron retention. This result strongly suggests that exon definition is a widely used physiological mechanism of SS recognition. Surprisingly, it is exons with the strongest 5’ SSs that are preferentially skipped when U1-70K is limiting, suggesting that U1-70K dependent activation of stable U1 snRNA-5’SS complexes is important for exon definition.

Complementation of the degron with transgenes showed that mutation of the RNAPII interaction surface of the U1-70K RRM causes extensive exon skipping compared to WT. These results suggest that U1 snRNP contact with the polymerase enables exon definition, possibly by favoring an optimal orientation for making cross-exon contacts with U2AF at the upstream 3’SS (Shao et al. 2025).

## Results

### U1-70K independent pausing by RNAPII at the first base of introns

We engineered HCT116 cells with a homozygous C-terminal *E. coli* dhfr degron at U1-70K which confers trimethoprim (TMP) dependent protein stabilization (Sheridan and Bentley 2016). Following TMP withdrawal, most U1-70K is degraded within 6hr and further depletion occurs after 24 hr (Fig. 1A). To investigate how U1-70K might influence transcription, we sequenced transcripts pulse labelled with bromouridine and purified by immunoprecipitation (Bru-seq) (Paulsen et al. 2014). In agreement with the effects of a U1 snRNA AMO (Mimoso and Adelman 2023), U1-70K depletion (22 hr) specifically reduced nascent RNA synthesis at 5’ ends of genes (Fig. 1B) consistent with a defect in RNAPII processivity. To investigate transcription at single nucleotide resolution we performed eNETseq which maps the 3’ ends of nascent transcripts co-immunoprecipitated with RNAPII (Churchman and Weissman 2011; Nojima et al. 2015). U1-70K depletion caused a subtle but reproducible narrowing of the promoter proximal peak of paused RNAPII consistent with impaired early elongation or increased premature termination (Fig. S1A)

**Figure 1.**
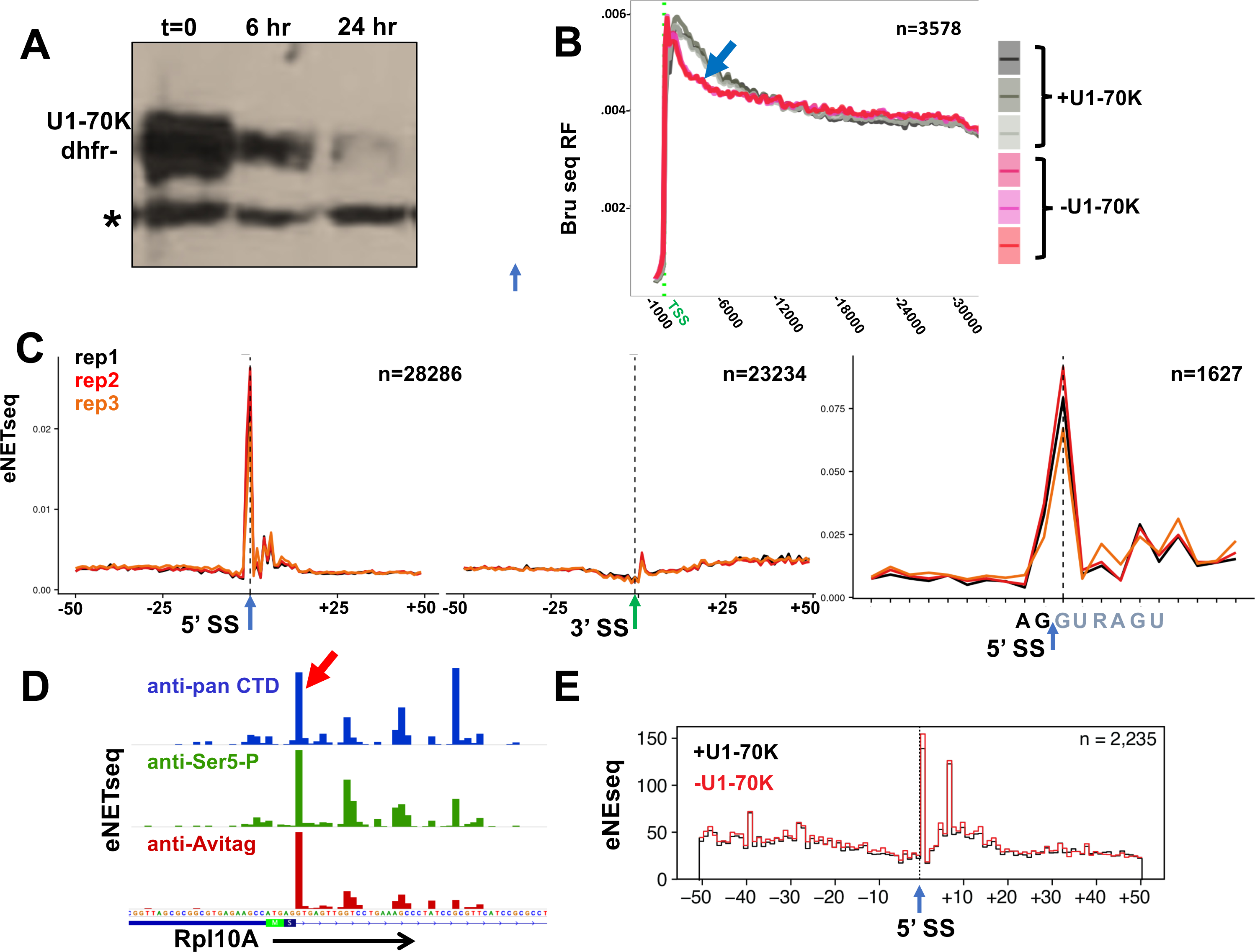
**A**. Degron depletion of U1-70K at time points after TMP washout. Western blot of HA-tagged dhfr degron fusion protein in HCT116 cells. * marks cross-reacting band. **B.** Reduced 5’ transcription elongation following U1-70K depletion. Nascent RNA sequencing relative frequency metaplots for replicate (n=3) Bru-seq experiments -/+ U1-70K degradation (36 hr) for genes >50kb long. Arrow marks signal drop suggesting reduced processivity at 5’ ends in U1-70K depleted cells. **C.** eNETseq metaplots (n=3) showing RNAPII pausing at the first base (+1,G) of introns in HEK293 Flp-in cells expressing Avitag WT Rpb1 Am^r^ (left and right panels) but not near 3’ SSs (middle panel). **D.** Screenshot of eNETseq in HCT116 (top track) HEK293 Flp-in cells (middle track) and HEK293 Flp-in expressing Avitag WT Rpb1 Am^r^ (bottom tracks) using three anti-RNAPII antibodies showing pausing at the first base (+1,G red arrow) of the first intron of RPL10A. **E.** eNETseq shows no effect of U1-70K degron depletion (22 hr) on pausing at the start of introns. A metaplot of first introns is shown.

RNAPII pauses are enriched at the G of G,T or G,C dinucleotides (Sheridan et al. 2019; Gajos et al. 2021; Fong et al. 2022). We therefore re-investigated whether pausing occurs at 5’ SSs and, if so, whether it is affected by U1 snRNP. We observed that eNETseq datasets generated by immunoprecipitation of Avitag epitope tagged Rpb1 in HEK293 Flp-in cells (Fong et al. 2022) had particularly low levels of contamination with step 1 splicing intermediates. These contaminants have 3’ ends at the 5’SS and confound the identification of pauses in the vicinity. By reanalyzing these datasets, we found that widespread pausing occurs after the first base of the intron at the canonical GT motif. We confirmed that pausing at the +1 base of introns occurs using two other anti-RNAPII antibodies (Fig. 1C, D). In contrast, a control anti Avitag eNETseq experiment in cells that do not express the epitope tagged RNAPII had a high level of 3’ ends at the 5’ SS from contaminating splicing intermediates (Fig. S1B). eNETseq in degron cells showed that U1-70K depletion did not noticeably affect pausing at the first base of the intron (Fig.1E, S1E). Whether pausing at this position, which precedes synthesis of the full 5’ SS sequence and extrusion from the RNA exit channel, influences co-transcriptional splicing remains to be determined.

### U1-70K degron depletion reveals widespread exon definition

We investigated how U1-70K depletion affected co-transcriptional splicing by sequencing of nascent transcripts co-immunoprecipitated with RNAPII that was solubilized by digestion of nuclei with DNase I (tNETseq) (Fong et al. 2017). This experiment which assays mostly constitutive co-transcriptional splicing events showed that overall splicing efficiency (SE) (Saldi et al. 2021) was only modestly reduced by U1-70K depletion (Fig. S1C), consistent with *in vitro* results using reconstituted U1 snRNPs lacking U1-70K (Will et al. 1996; Rogalska et al. 2024).

We performed polyA+ RNA-seq in U1-70K degron cells and examined alternative splicing. This experiment demonstrated a remarkably high level of exon skipping and much lower levels of exon inclusion and intron retention when U1-70K was depleted for either 8hr or 36 hr (Fig. S1D, 2A). After 36 hr of U1-70K depletion, we detected over 6500 examples of increased alternative exon skipping and fewer than 500 examples of intron retention (Fig. 2A-C, S2A-C). In summary, the overwhelming response to disruption of U1 snRNP function by U1-70K depletion is exon skipping, which is the predicted result if exon definition is a major mechanism of SS recognition in the human transcriptome. This conclusion is supported by the observation that exons skipped in response to U1-70K depletion are shorter than unaffected or included exons and are flanked by longer introns (Fig. 3A) which both favor exon definition (Fox-Walsh et al. 2005; Hollander et al. 2016; Pai et al. 2017; Carranza et al. 2022). Transcriptional pausing was observed at the 5’ SSs of skipped exons using NET-seq, but it was not significantly affected by U1-70K depletion (Fig. S3E).

**Figure 2.**
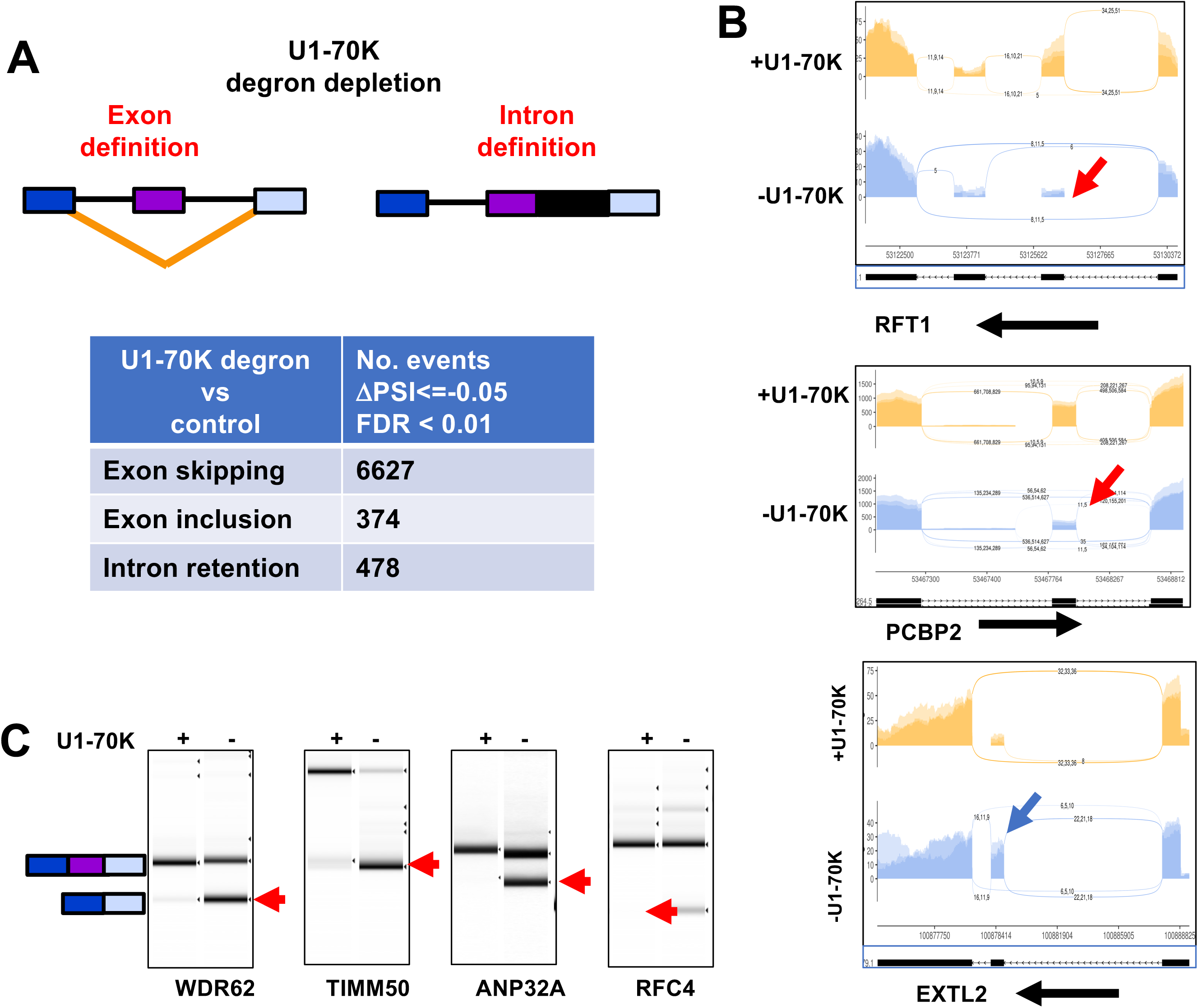
**A**. PolyA+ RNA-seq (n=3) datasetd before and after U1-70K degron depletion (36hr) were analyzed by rMATS (Shen et al. 2014) revealing widespread exon skipping consistent with exon definition. **B.** Screenshots made with ggSashimi (Garrido-Martín et al. 2018) for 3 replicate RNA-seq datasets showing exon skipping (red arrows) and inclusion (blue arrow) with U1-70K depletion. **C.** RT-PCR confirmation of several examples of exon skipping detected by RNA-seq following U1-70K depletion.

**Figure 3.**
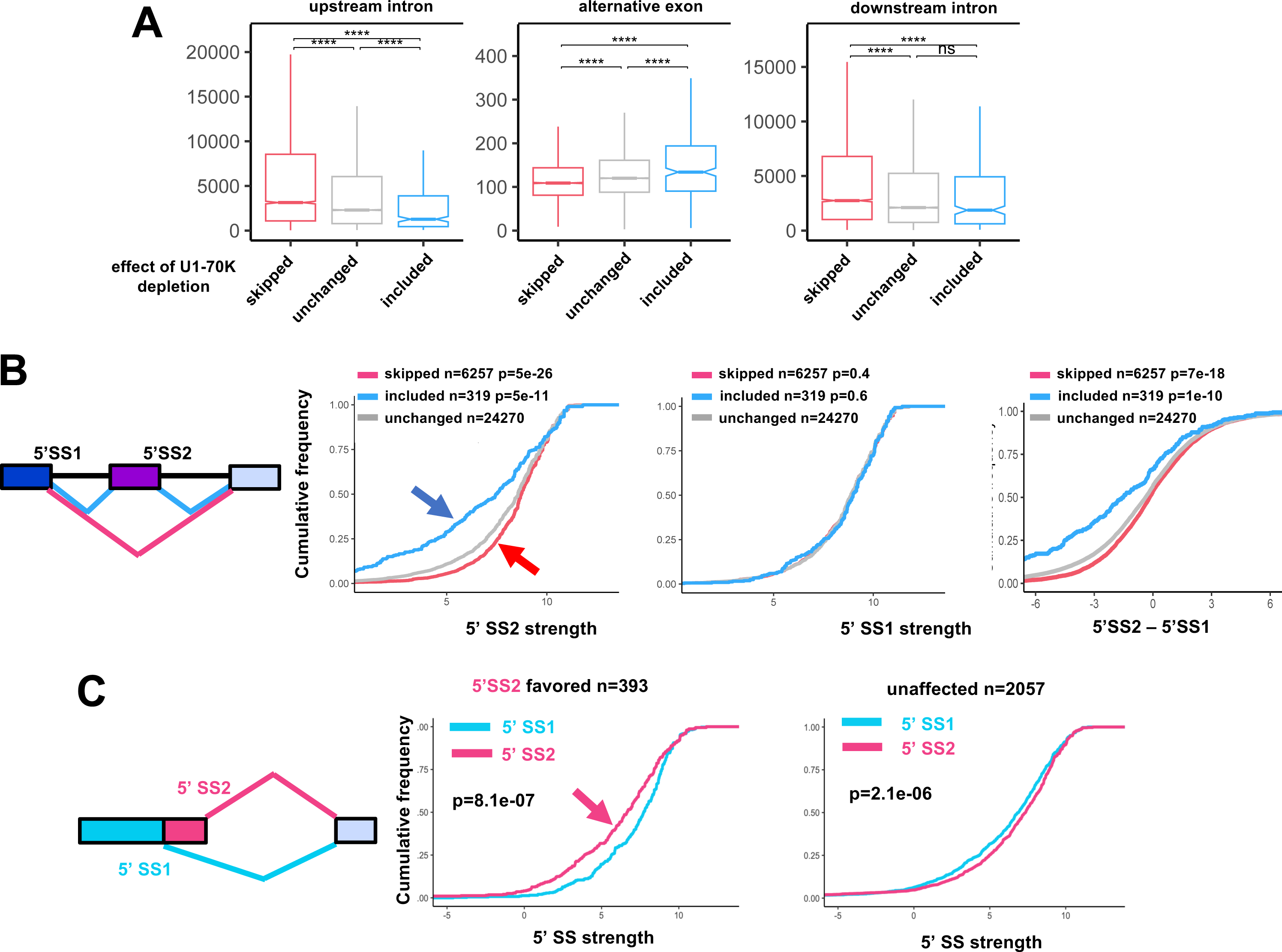
**A**. Properties of unaffected, skipped, and included exons following U1-70K degron depletion. Note skipped exons have longer upstream and downstream flanking introns and are shorter in length than unaffected or included exons as expected if they are recognized by exon definition (****p ≤ 0.0001 Wilcoxon Rank Sum test). **B.** Cumulative frequency plots of 5’SS strengths determined by MaxENT for 5’SS1 and 5’SS2. Note significantly stronger 5’SS2 at exons skipped (pink) and weaker 5’SS2 at exons included (blue) with U1-70K depletion (left panel) but no differences in strength of 5’SS1 (middle panel). Right panel shows the relative strengths of adjacent pairs of 5’SS1 and 5’SS2. Note 5’SS2 is stronger than 5’SS1 when the exon is skipped in response to U1-70K loss. **C.** Cumulative frequency plots as in B for alternative 5’SSs. Note that U1-70K depletion favored used of the weaker downstream site (5’SS2, left panel) in contrast to unaffected alternative 5’SSs (right panel).

U1-70K depletion, in contrast to U1 AMO treatment, did not strongly activate premature cleavage polyadenylation (PCPA) within introns (Kaida et al. 2010; Feng et al. 2023) (Fig S3A-C). This result is consistent with the small effect of the U1-70K degron on overall splicing efficiency (Fig. S1C), and with the hypothesis that PCPA results from reduced competition with intron removal by splicing (Peterson and Perry 1989; Feng et al. 2025).

### U1-70K depletion causes skipping of exons with strong 5’ splice sites

We compared the strengths of 5’ SS’s (Yeo and Burge 2004) for skipped exons with those that are unaffected by U1-70K depletion. Surprisingly, the skipped exons have 5’ SSs that are significantly *stronger* than unaffected exons. Conversely, the 5’ SSs of the minority class of exons that are more included when U1-70K is depleted, are *weaker* than at other exons (Fig. 3B, left panel). On the other hand, the strengths of the 5’SS upstream of skipped exons (5’SS-1) were not significantly different from unaffected or included exons (Fig. 3B middle panel). If exon definition is required for splice site recognition, then the decision between inclusion versus skipping of an exon can be viewed as a competition between upstream (5’SS1) and downstream (5’SS2) 5’ SSs where inclusion results if the downstream 5’ SS prevails in making a productive cross-exon complex. We therefore compared the strengths of individual pairs of 5’ SSs at skipped exons and their upstream constitutive exons. Notably, the 5’ SSs of exons skipped under U1-70K limiting conditions are generally stronger than the competing upstream sites, but this is not the case for unaffected or included exons (Fig. 3B, right panel).

We investigated whether the bias against use of strong 5’ SSs when U1-70K is limiting also applied to the competition between alternative 5’ SSs (A5’SSs). Almost 400 examples were identified where U1-70K depletion favored a downstream A5’SS. Remarkably A5’SSs favored for splicing under these conditions were significantly weaker than the competing upstream sites, in contrast to unaffected pairs of A5’ SSs (Fig. 3C), mimicking the results seen with skipped exons. In summary, under U1-70K limiting conditions, weak 5’ SSs are paradoxically more successful in supporting productive splicing than stronger 5’SSs, both at skipped exons and alternative 5’ SSs.

### Poison exons are frequently included in response to U1-70K depletion

To ensure that nonsense mediated decay of mis-spliced transcripts did not bias our results, we repeated RNAseq in U1-70K degron cells in the presence of the NMD inhibitor SMGi (Gopalsamy et al. 2012). The experiment confirmed that U1-70K degradation causes pervasive exon skipping (Fig. 4A). An exception is the group of ultra-conserved nonsense codon-containing poison exons (Lareau et al. 2007) that are characteristic of SR protein genes and are stabilized when NMD is inhibited. These exons including those in SRSF2, 3, 7 and SMN2 (exon7a) and SRRM1 are more included in response to U1-70K depletion (Fig. 4B, C, S2D-F). Notably, poison exons have weak downstream 5’ SSs relative to exons unaffected by the U1-70K degron (Fig. 4D) lending further support to the notion that limiting U1-70K specifically favors inclusion of alternative exons with weak 5’SSs. Whether poison exon inclusion under U1-70K limiting conditions serves a physiological function remains to be investigated.

**Figure 4.**
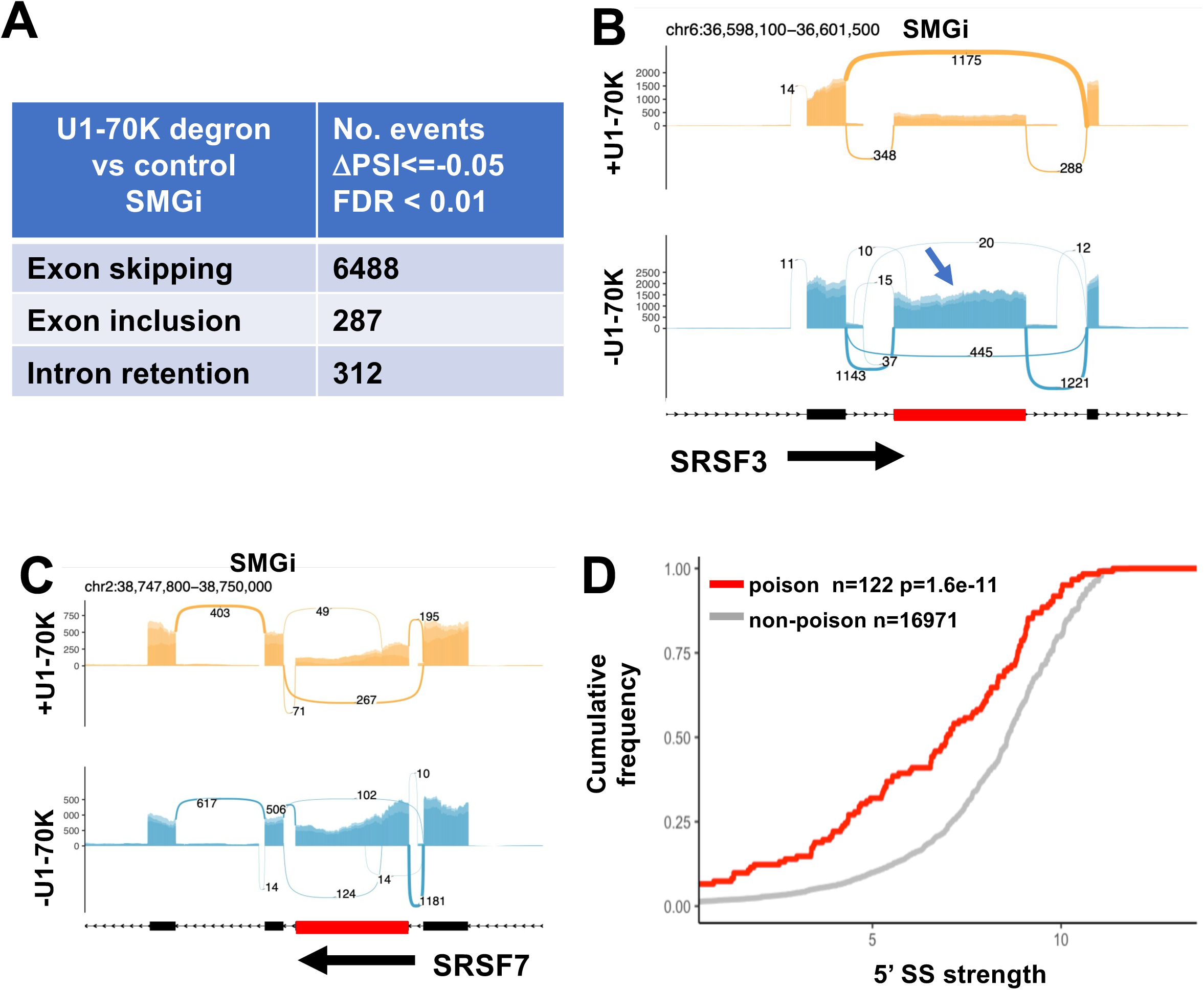
**A**. RNA-seq (n=3) datasets of rRNA depleted total RNA before and after U1-70K degron depletion (36hr) in the presence of the NMD inhibitor SMGi (0.5μM 24 hr) were analyzed by rMATS (Shen et al. 2014) revealing widespread exon skipping consistent with exon definition. **B. C.** Screenshots made with ggSashimi (Garrido-Martín et al. 2018) for 3 replicate RNA-seq datasets showing enhanced inclusion (blue arrows) of poison exons (red boxes) with U1-70K depletion. **D.** Poison exons (Thomas et al. 2020) have weaker 5’SSs than other exons. Cumulative frequency plots of 5’SS strength as in Fig. 3B.

### Strengthening of 5’splice sites induces exon skipping when U1-70K is limiting

If 5’ SS strength determines susceptibility to skipping, then strengthening a SS is predicted to induce more skipping of the corresponding exon when U1-70K is limiting. We identified three exons in GOLGB, DLGAP5 and CEP350 that have different non-canonical 5’SS sequences whose inclusion is relatively unaffected by U1-70K depletion and expressed them in CMV driven minigenes with their flanking exons (Fig. 5A). The minigenes were transiently transfected into U1-70K degron cells with expression vectors for WT U1 snRNA or a mutant snRNA that fully complemented the non-canonical 9 base 5’ SS sequence of each exon (Fig. S3E). Exon inclusion was assayed by RT-PCR in cells before and after U1-70K degradation. In each case, exon skipping induced by U1-70K depletion was enhanced by the cognate mutant U1 snRNA that conferred full base pairing with the 5’ SS relative to the mismatched WT U1 (Fig. 5A).

**Figure 5.**
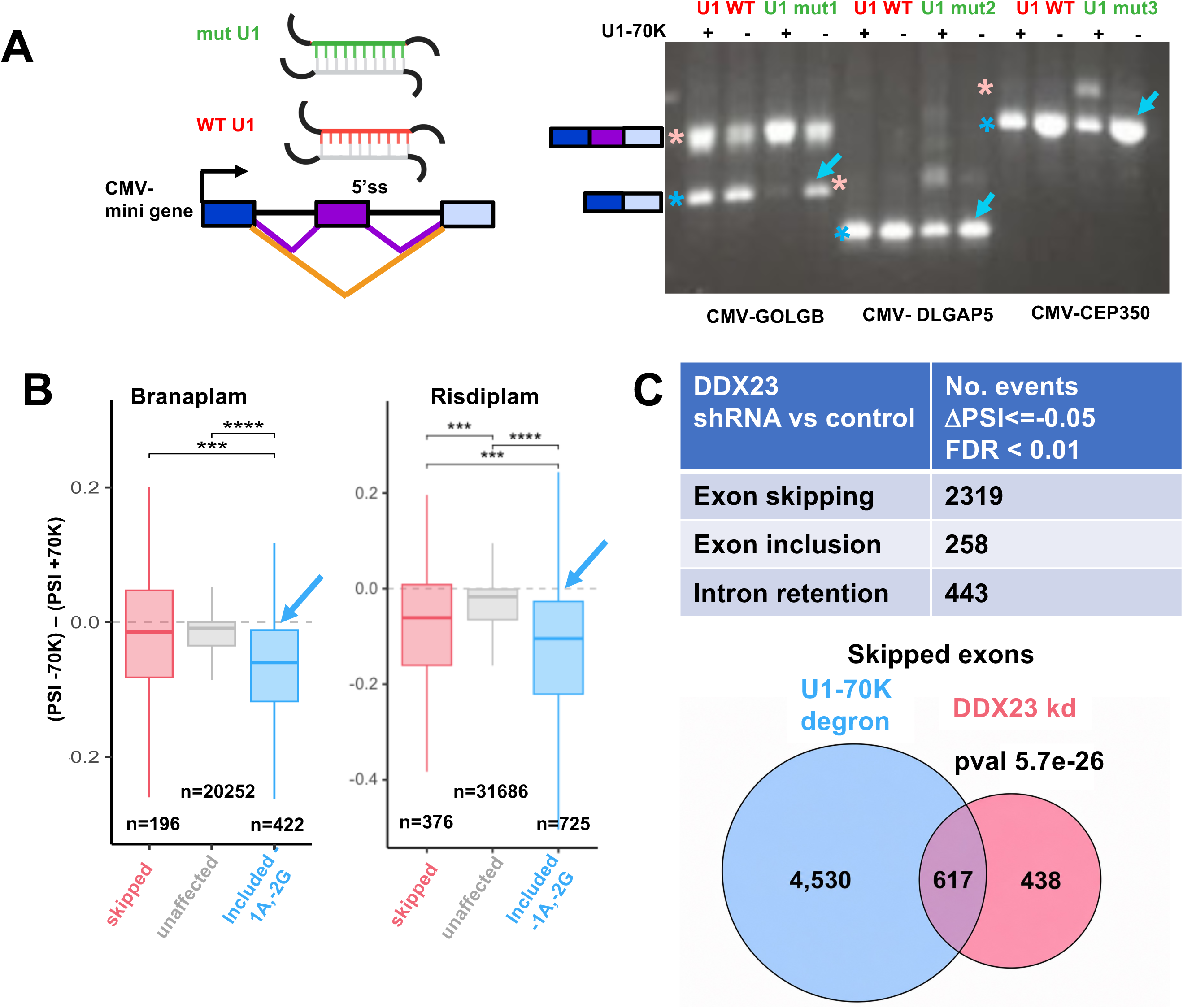
**A**. Skipping of alternative exons is specifically enhanced by U1-70K depletion when their 5’SSs are strengthened with complementary U1 snRNAs. CMV driven minigenes contained alternative exons and flanking regions from GOLGB (exon 7), DLGAP5 (exon 19) and CEP350 (exon 23) that were transiently transfected into U1-70K degron cells along with expression vectors for WT and cognate mutant U1 snRNAs (Fig. S3E) that restored full base pairing to the respective 5’ SSs. RT-PCR (right panel) shows that the mutant U1snRNAs confer enhanced skipping in response to U1-70K depletion (blue arrows). Pink and blue * to the left of each panel denote RT-PCR products for exon inclusion and skipping respectively. **B.** Splice modifiers branaplam and risdiplam confer enhanced skipping of target exons when U1-70K is depleted. We identified variant exons with -1A, -2G 5’SSs that more included or more skipped in response to branaplam or risdiplam in HCT116 U1-70K degron cells under control conditions (+TMP). These exons were monitored for the effect of U1-70K depletion (36hr) ((PSI -70K) - (PSI +70K)) in the presence of the drug (36hr. 70nM branaplam, 140 nM risdiplam). Note that branaplam and risdiplam specifically enhance skipping of sensitive exons in response to U1-70 depletion (blue arrows). (***p ≤ 0.001, ****p ≤ 0.0001 Wilcoxon Rank Sum test). **C.** DDX23/PRP28 kd mimics U1-70K depletion effect on exon skipping. rMATS analysis of RNAseq (n=3) from A2780 ovarian cells treated with control or DDX23 siRNA (Zhao et al. 2021). Lower panel: significant overlap between exons skipped in U1-70K depleted HCT116 and DDX23 kd A2780 cells. Only exons detected in both datasets are included.

Non-canonical 5’ SSs characterized by A at -1 and G at -2 relative to the splice junction can be strengthened by the RNA targeting drugs branaplam and risdiplam (Campagne et al. 2019; Ishigami et al. 2024; White et al. 2024; Kuang et al. 2026). If SS strength governs exon skipping in U1-70K limiting conditions, then skipping of -1A, -2G exons should be specifically enhanced by these drugs when U1-70K is degraded. Several hundred exons with -1A -2G 5’ SSs were identified that are more included in the presence of branaplam or risdiplam in cells with wildtype U1-70K levels (+TMP).

We then asked how inclusion of these drug-sensitive exons was affected by U1-70K depletion compared to unaffected exons. In the absence of the splice modifier, sensitive -1A, -2G exons were not affected by U1-70K depletion more than other exons. In contrast, in the presence of branaplam or risdiplam -1A, -2G exons were significantly more skipped than other exons when U1-70K was depleted (Fig. 5B). In summary, strengthening a 5’SS either with a complementary mutant U1 snRNA or with a small molecule confers higher levels of skipping when U1-70K is depleted. We conclude that 5’SS strength is a critical determinant of whether an exon is skipped under U1-70K limiting conditions and that this U1 snRNP subunit is specifically required for splicing at strong 5’ SSs.

One way to rationalize the paradox that strong 5’ SSs specifically inhibit splicing when U1-70K is limiting is that under these conditions they form unproductive hyper-stable complexes with U1 that cannot efficiently undergo DDX23/Prp28 mediated transfer to U6. This model predicts that DDX23 knock down might mimic the U1-70K degron and enhance exon skipping. We tested this idea by examining alternative splicing in published RNA-seq data sets for DDX23 knock down in human ovarian cells (Zhao et al. 2021) and CRISPR mediated knock out in K562 cells (Consortium 2012). In both studies, the major effect of DDX23 depletion was to promote exon skipping (Fig. 5C, S4A). Thus, knockdown of the DDX23 helicase mimics U1-70K depletion, supporting the idea that these two proteins cooperate to activate stable U1-5’SS complexes for exchange with U6 and facilitate the definition of exons with strong 5’ SSs.

### The RNAPII interaction surface of U1-70K is important for exon definition

To investigate the role of contact between U1-70K and RNAPII in co-transcriptional splicing we mutatedg residues in U1-70K that are closest to the Rpb2 and Rpb12 subunits of RNAPII in the complex with U1 snRNP (Zhang et al. 2021). U1-70K degron cells were infected with lentiviral vectors enabling doxycycline inducible expression of Avitag epitope tagged WT U1-70K or a mutant (mut7) with 7 substitutions in the RNAPII interaction surface in α helices 1 and 2 of the RRM (K118E, R121E, E124K, V125A, Y126A, E152K, D156K, Fig. 6A). The mut7 protein was expressed at similar levels to the WT (Fig. 6C, lanes 1, 6). We compared RNAPII binding of U1 snRNP bearing U1-70Kmut7 with WT by pulling out RNAPII complexes on GST-TFIIS beads in an assay previously used to detect this association (Robert et al. 2002) (Fig. 6B). TFIIS binds to RNAPII by inserting a Zn finger domain into the side channel of the polymerase. Complexes eluted from the GST-TFIIS beads were assayed by western blotting to detect Avitag U1-70K WT or mut7 as well as endogenous U1A. Relative to WT U1-70K, a smaller fraction of the mut7 protein associated with RNAPII compared to input (Figure 6C, compare lanes 1 and 3 with 6 and 8) suggesting that mutation of the Rpb2/Rpb12 interaction surface reduces, but does not abolish, binding of the snRNP to RNAPII. This result is consistent with the finding that a triple mutant (K118E, R121E, E124K) reduces binding of truncated purified U1-70K with RNAPII (Li et al. 2026). Residual binding by U1-70K mut7 U1 snRNP to RNAPII could be through binding to nascent transcripts as the extract was not RNAse digested in order to preserve snRNP integrity.

**Figure 6.**
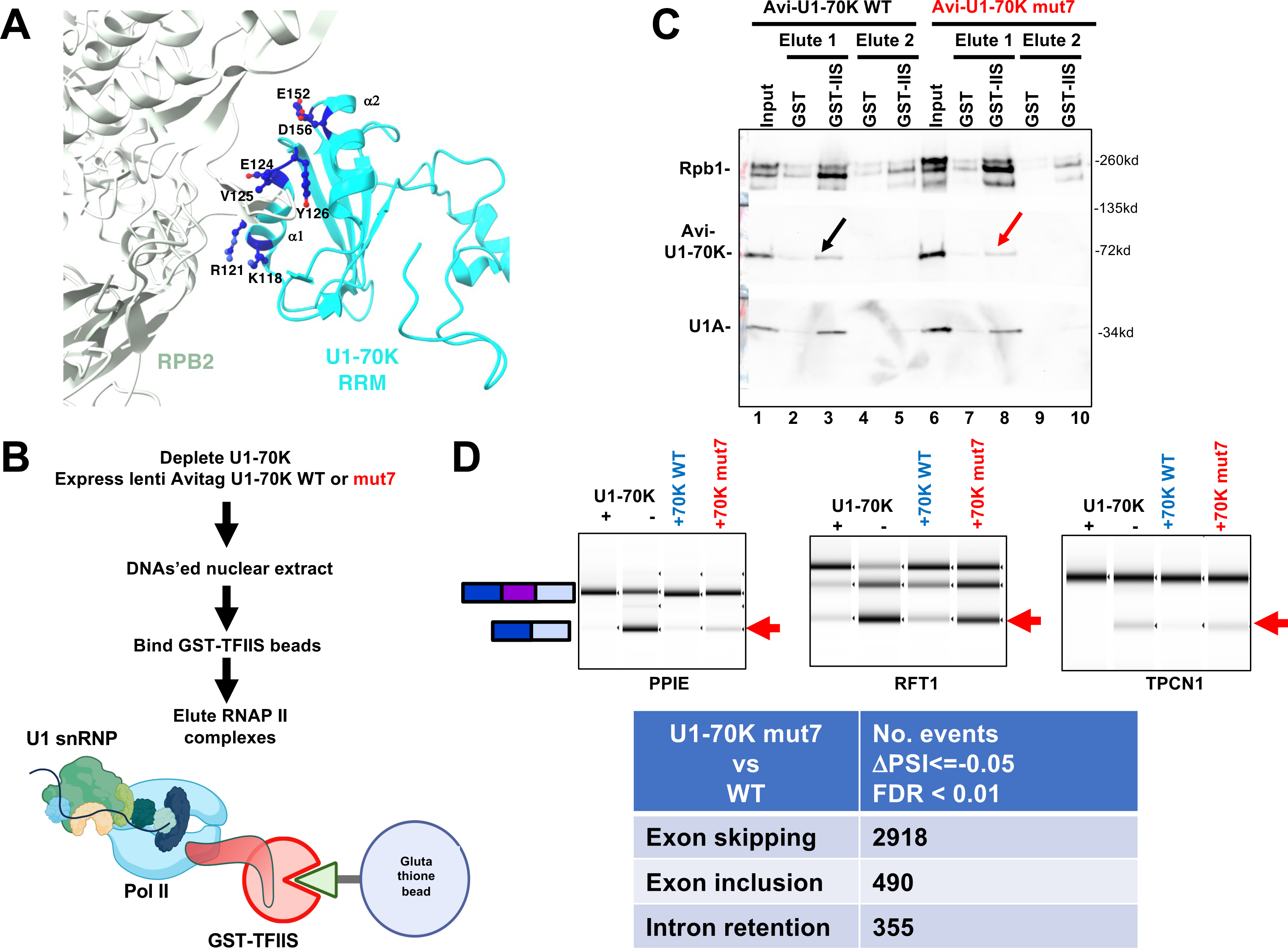
**A**. Model of the U1-70K RRM-RPB2 interface showing 7 residues in closest proximity to RNAPII (7B0Y)(Zhang et al. 2021) in alpha helices α1 and α2 that were mutated in the mut7 construct. **B.** Strategy for capturing RNAPII-U1-snRNP complexes on GST-TFIIS beads (Robert et al. 2002) from extracts of U1-70K degron cells complemented with WT or mut7 Avitag-U1-70K expressed from lentiviruses. **C.** U1-70K mut7 reduces U1 snRNP binding to RNAPII. Successive high salt eluates from GST-TFIIS beads from B were probed by Western blotting for Rpb1, Avitag-U1-70K, and U1A. Note that relative to eluted Rpb1, mut7 binding is modestly reduced relative to WT U1-70K. **D.** U1-70K mut7 causes widespread exon skipping relative to WT U1-70K. polyA+ RNAseq (n=3) from U1-70K depleted cells (36hr) complemented with lentiviral expressed (+ doxycycline 24 hr) WT or mut7 U1-70K. Lower panel rMATS analysis as in Fig. 2A. Upper panel RT-PCR validation of exon skipping specific to mut7 (red arrows).

We compared splicing in cells where U1-70K degron depletion was complemented by exogenous WT or mut7 U1-70K. WT U1-70K fully rescued the exon skipping phenotype of the degron. Whereas U1-70K mut7 restored inclusion of many skipped exons, we identified almost 3000 high confidence examples of exon skipping that persisted in mut7 expressing cells relative to WT (Fig 6D). Like exons skipped in response to U1-70K depletion, those skipped with the U1-70K mut7 mutant had stronger 5’SSs, and those that were included had weaker 5’SSs than unaffected exons (Fig. S4B). There was extensive and highly significant overlap between the exons skipped in response to U1-70K depletion and mutation of the RNAPII interaction surface (Fig. S4C). We conclude that perturbing the RNAPII interaction surface of the U1-70K RRM disrupts definition of a large subset of exons.

## Discussion

In this report we used controlled degron degradation of the U1-70K subunit of U1 snRNP to investigate the *in vivo* importance of exon definition and to query the role of direct U1 snRNP contact with RNAPII in this process. The in vivo relevance of exon definition has been brought into question by the discovery ultra-fast splicing that occurs before transcription of a downstream exon is completed (Oesterreich et al. 2016; Zeng et al. 2022). Depleting cells of U1-70K is expected to prevent direct contact between U1 snRNP and the polymerase and to disrupt 5’SS recognition which in turn will cause skipping if exon definition operates (Robberson et al. 1990). The salient conclusions from this work are:

1. U1-70K depletion inhibited transcription elongation in 5’ regions of genes consistent with previous results using a U1 AMO (Mimoso and Adelman 2023). We report that RNAPII pausing occurs after the first base of introns at the canonical GT motif but it is not affected by U1-70K depletion and is probably determined predominantly by DNA sequence.
2. U1-70K depletion caused skipping of thousands of exons strongly suggesting that exon definition operates widely in vivo consistent with the effects of 5’SS mutants (Krawczak et al. 2007) and that this process specifically requires U1-70K in agreement with previous work (Rogalska et al. 2024).
3. U1-70K deficiency specifically favors use of weak 5’ SSs and disfavors use of strong 5’ SSs both at alternative cassette exons and alternative 5’SSs (Fig. 3B, C). Furthermore, strengthening of 5’ SSs sensitizes exons to skipping when U1-70K is limiting (Fig. 5). These results therefore suggest that U1-70K facilitates formation of productive exon definition complexes with strong 5’SSs, consistent with our observation that knockdown of the DDX23/Prp28 helicase that exchanges U1 for U6 snRNA also causes mostly exon skipping (Fig. 5C, S4A).
4. The Cryo-EM structure of the human U1 snRNP-RNAPII complex indicates that U1 snRNP cannot make direct contact with RNAPII in the absence of U1-70K (Zhang et al. 2021). Therefore, the effect of depleting this subunit on exon definition is consistent with the scenario that contact with the polymerase enables formation of cross-exon complexes as previously suggested (Shenasa and Bentley 2023; Shao et al. 2025). This idea is further supported by our finding that the major effect of mutating 7 residues in the U1-70K RRM at the interface with Rpb2/12 is to cause exon skipping (Fig. 6D) though it is less extensive than that caused by U1-70K depletion.

Transcriptional pausing at or near SSs has been proposed to help coordinate transcription with co-transcriptional pausing (Alexander et al. 2010; Milligan et al. 2017) but it has been difficult to unambiguously distinguish paused nascent transcripts from splicing intermediates that co-purify with RNAPII. We report that in NET-seq datasets free of significant contamination with splicing intermediates, preferential RNAPII pausing is revealed at the first G of introns (Fig. 1C, D). This observation is consistent with the known preference for pausing at G’s that precede a pyrimidine (Fong et al. 2022). Whether pausing at the first base of the intron, which is 15-20 bases before the 5’SS emerges from the RNA exit channel, influences co-transcriptional splice site recognition remains unknown, however pausing at this position was not significantly affected by U1-70K depletion.

Two questions raised by these findings are: "Why are strong 5’SSs selected against when U1-70K is limiting?" and "How is the strength of a 5’SS at a skipped exon sensed when it does not actually participate in productive splicing?". We suggest that U1 snRNP engages the 5’SS of the downstream exon prior to splicing of the upstream intron as proposed for classical exon definition. If the 5’ SS is strong, and if U1-70K is absent, then U1 snRNP is hyper-stabilized and this impairs the transition from U1 to U6 engagement and ultimately results in skipping of the exon. This model is consistent with the fact that yeast U1 snRNP interacts much more dynamically with the 5’SS than is expected based solely on RNA base-pairing interactions (Hansen et al. 2022). We note that the model of U1 binding and stabilization at downstream exons with strong 5’SSs prior to splicing of the upstream intron, is not compatible with a U1 relay (Yoon et al. 2026) functioning at these exons.

Extended base pairing of human U1 snRNA with the 5’SS to make hyper-stable complexes can favor exon inclusion (Freund et al. 2005). We speculate that splicing under these conditions likely requires U1-70K to facilitate DDX23/Prp28 mediated exchange of U1 for U6 snRNA. On the other hand, exons with weak 5’SSs that make less stable U1 snRNA-5’SS complexes, are preferentially included when U1-70K is limiting (Fig, 3B). This observation suggests that U1-70K normally suppresses inclusion of such exons possibly by impairing formation of sufficiently stable complexes with U1 snRNP at their 5’SSs.

The effects of U1-70K limitation, strongly suggest that exon definition is a widely used mechanism of SS recognition *in vivo*. The importance of the U1-70K subunit, which contacts RNAPII, and in particular its RNAPII interaction surface, for exon definition indicate that this process frequently occurs on transcribing RNAPII. Specifically, our results are consistent with the model that exon definition is initiated by interaction of U1-snRNP engaged with a 5’SS, and U2AF engaged with a 3’SS, while they are co-localized on the RNAPII surface (Shenasa and Bentley 2023; Shao et al. 2025) (Fig. 7). We cannot exclude the possibility that the U1-70K RRM surface that contacts Rpb2/12 also interacts with other partners that contribute to exon definition. Notably SRSF1 promotes exon definition and is the best characterized interactor with the U1-70K RRM (Wu and Maniatis 1993; Cho et al. 2011). However SRSF1 interaction is unlikely to mediate all the effects we observed because the C-terminal extension of the RRM required for SRSF1 interaction (Paul et al. 2024) does not overlap with the RNAPII interaction surface that we mutated (mut7, Fig. 6A) on the α1 and α2 helices (Zhang et al. 2021). We speculate that weaker binding and/or a non-optimal orientation of U1 snRNP on the RNAPII surface in the absence of U1-70K (Fig. 7) impairs some aspect of exon-defined spliceosome maturation (Zhang et al. 2024a). One possibility is that U1-70K helps recruit or orient DDX23/Prp28 in the pre-B complex together with SRSF1 (Cho et al. 2011; Segovia et al. 2024) or other intermediary proteins thereby facilitating U1-U6 exchange and the pre-B to B complex transition that results in exon inclusion. In the absence of U1-70K hyper-stable U1-5’SS interaction may lead to formation of non-productive complexes, possibly analogous to those formed in response to negative regulators of splicing (House and Lynch 2006; Sharma et al. 2008), that ultimately result in exon skipping. In future it will be important to elucidate at the atomic level how U1-70K functions in the pre-B to B complex transition in the context of a transcription complex.

**Figure 7.**
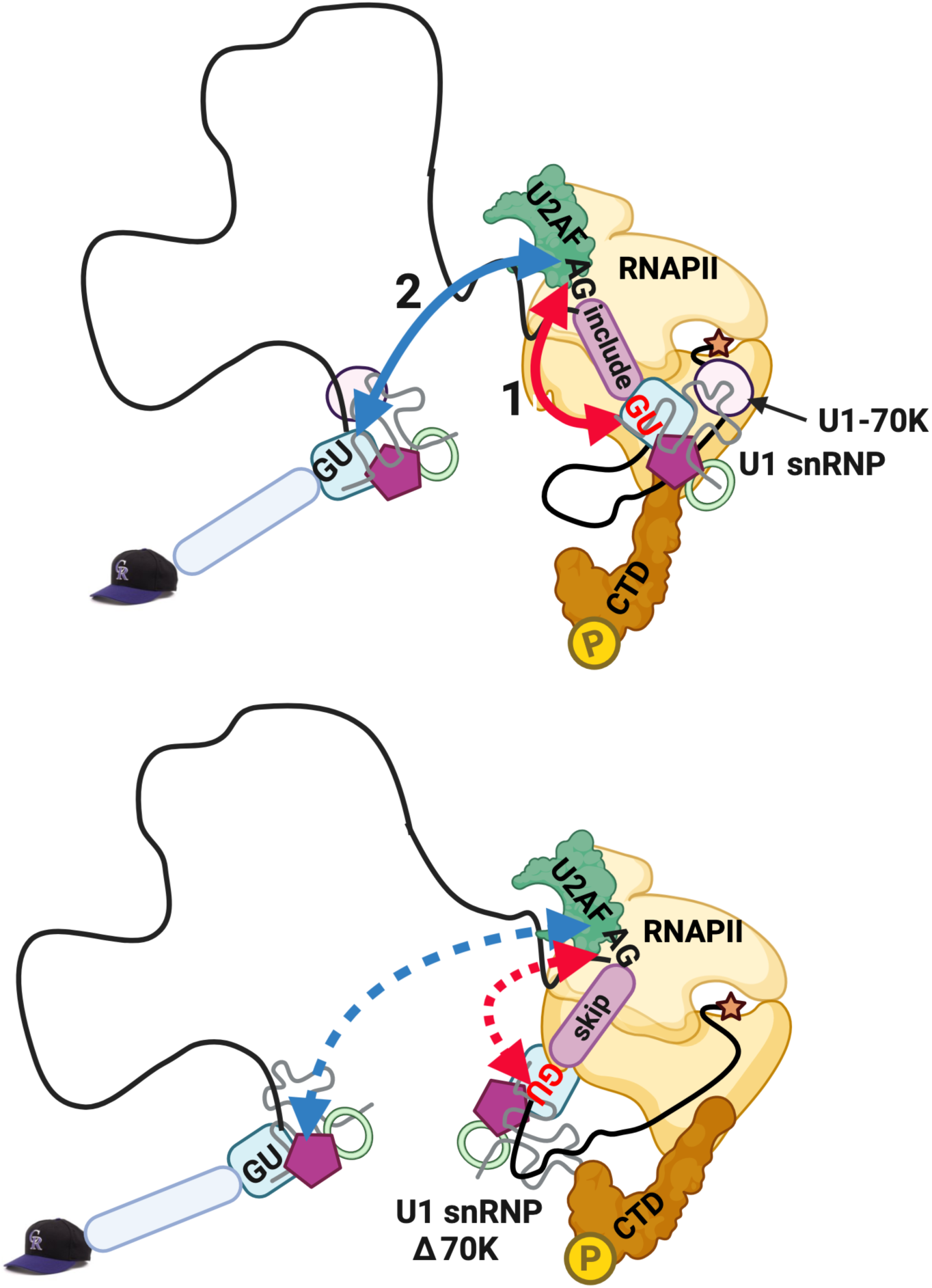
Model of co-transcriptional exon definition facilitated by U1-70K interaction with RNAPII (top panel) and its disruption in the absence of U1-70K (bottom panel). Cross-exon (red arrow, 1) followed by cross-intron (blue arrow, 2) complex formation (top panel) resulting in inclusion of the downstream exon (purple) is impaired by loss of U1-70K (bottom panel) resulting in skipping. GU and AG motifs at the 5’SS and 3’SS engaged by U1 snRNP and U2AF are shown. Star signifies 3’ end of nascent transcript. Made with Biorender.

## Materials and Methods

### Human cell lines and cell culture

A C-terminal E coli dhfr degron was inserted at both copies of U1-70K in HCT116 cells with an HA epitope tag and in-frame T2A-neo or T2A-hygro selectable markers as described (Sheridan and Bentley 2016). Cells were maintained in McCoy’s medium supplemented with 10% FBS and penicillin/streptomycin plus 200 μg/ml hygromycin B, 1 mg/ml G418 and 10μM trimethoprim (TMP). The degron was activated by washing out TMP.

HEK293 Flp-in (Invitrogen) cells expressing Avitag Am^r^ Rpb1 WT (Fong et al. 2022) were maintained in DMEM 10% FBS, 200 μg/mL hygromycin B, 6.5 μg/mL blasticidin, 1% pen/strep.

For RNA-seq in U1-70K degron cells RNA was harvested with Trizol 8hr or 36 hr after TMP washout. SMGi treatment was for 24 hr at 0.5 μM, branaplam was for 36 hr at 70 nM and risdiplam was for 36 hr at 140 nM (Ishigami et al. 2024).

### Transient Transfection

10 cm plates of U1-70 degron cells were transfected with 2.5μg or pMT14 minigene plus 2.5 μg of pAT U1 snRNA expression vector using Lipofectamine 2000 (10 μg) and RNA was harvested after 48 hr. with Trizol.

### Lentiviruses

U1-70K degron cells were infected with lentiviruses based on pCW57 a gift of Adam Karpf (Addgene 89180) and selected with blasticidin (10 μg/ml). The lentiviruses expressed codon optimized WT and mutant U170K ORFs with a C-terminal AvitagX2 that were synthesized by Genewiz. Doxycycline induction (2 µg/mL) was for ∼24 hr.

### Plasmids

Minigenes for GOLGB1, CEP350 and DLGAP5 alternate exons with non-canonical 5’SSs were constructed in the pMT14 vector by Synbio Technologies with CMV promoter, alpha globin 5’UTR, initiation codon with Kozak consensus, termination codon, and HSV TK polyA site. Sequences of the inserts are available on request.

U1 snRNA expression vectors pAT U1 WT, mut1, 2, 3 were constructed with inserts synthesized by Twist Bioscience and cloned between the Bgl II and Hind III sites of the human U1 snRNA plasmid HU1-1D in pAT153 (Neuman de Vegvar et al. 1986).

GST-TFIIS was expressed in E. coli from pGEX2T TFIIS in Topp2 cells (Pan et al. 1997).

**RT-PCR** used cDNA from DNAse I treated total RNA primed with random hexamers using M-MLV RT. Primers are provided in Table S1. RT-PCR products were visualized with a Bioanalyzer.

### Antibodies

Homemade rabbit anti-RNAPII pan CTD (Schroeder et al. 2000) and anti-Avitag (Fong et al. 2022) antibodies have been described. Commercial antibodies: Mouse monoclonal anti HA 12CA5; rat monoclonal anti RNAPII CTD Ser5-P 3E8 (Chromotek); mouse anti BrdU 3D4 (Pharmingen 555627) and anti U1A (Abcam AB155054).

### GST-TFIIS purification of RNAPII complexes

Cells from two 15 cm plates of HCT116 U1-70K degron cells were lysed in 300mM sucrose,15mM Tris HCl pH8.0,15mM NaCl,60mM KCl, 1mMEDTA, 0.5mM EGTA, 0.15 mM Spermine, 0.5 mM spermidine, 0.5 mM DTT, 0.05% NP40 + protease inhibitors (He et al. 2014) 15 min on ice, then nuclei were pelleted, washed with 1 ml DNAse I buffer (1mM MnCl2, 10 mM NaCl, 1 mM CaCl2, 40mM Tris-HCl 7.9, 0.3 mM Spermine, 0.5 mM spermidine) and the pellet resuspended in 20 X volume DNAse I buffer + 33U/ml DNAse I (NEB #0303). DNAse I digestion was for 15 min at RT followed by 60 min at 4°C on a nutator. Nuclei were then lysed by sonication (10 min) and the extract cleared by centrifugation at 1500 rpm 5 min in a microfuge and supplemented with NaCl to 50mM, NP40 to 0.5% and glycerol to 8%. The extract was incubated with magnetic GST-TFIIS beads for 2 hr at 4° on a nutator. Beads were washed 2X in 1 ml 20mM Hepes; 10% Glycerol; 50mM NaCl; 1.5mM MgCl2 + 0.05% NP40 and eluted 2X in 125 μl of 20mM Hepes; 10% Glycerol; 300mM NaCl; 1mM EDTA; 0.05% NP40.

### Immunoblotting

Immunoblots were developed with HRP conjugated swine anti-rabbit secondary antibody (DAKO, P0217) and ECL Plus (Perkin Elmer, NEL103E001EA).

### RNA-seq

PolyA+ RNA or total RNA depleted of rRNA (Baldwin et al. 2021) was used to prepare RNA-seq libraries with the KAPA RNA HyperPrep kit (KR1350 – v1.16). Sequencing was on an Illumina NovaSeq 6000 (2x150). PCR duplicates were removed using bbtools clumpify and adapters were trimmed using bbtools bbkuk version 39.01. After filtering out rRNA, reads were mapped to the hg38 UCSC human genome with Bowtie2 version 2.3.2. Alternative splicing analysis was performed with rMATS version 4.0.2 (Shen et al. 2014). Alternative splicing events included in the analyses are supported by >50 reads in each of 3 replicates. Splice site strengths were determined with MaxENT (Yeo and Burge 2004). All significance values were calculated using the Wilcoxon Rank Sum Test.

### Bru-Seq

Nascent RNA sequencing by Bru-Seq was performed as described (Paulsen et al. 2014; Sheridan et al. 2019) with minor modifications. U1-70K degron cells + and – TMP (36hrs) were incubated with 2 mM Bromouridine in fresh medium for 30 min. Labeled RNA (50 μg) was fragmented with ZnCl2 (10mM, 70° 8 min, stopped with 10 mM EDTA) and immunopurified in PBS, 0.05% Triton X-100, 1 mM DTT, + RNase inhibitor for 1 hr at 4° using 3D4 monoclonal anti-BrdU (3 μg) immobilized on protein G Dynabeads. Beads were washed 3X in the same buffer and RNA was eluted in 5mM DTT at 95° for 3 min. Eluted RNA was rRNA depleted (Baldwin et al. 2021) and RNA-seq libraries were made with the KAPA RNA HyperPrep kit (KR1350 – v1.16). Mapping was as described for RNA-seq and reads were normalized to total counts on the mitochondrial chromosome.

### tNET-seq

tNETseq analysis of nascent transcripts co-immunoprecipitated with RNAPII using rabbit anti-pan CTD antibody from DNAse I treated nuclear extracts was performed as described (Fong et al. 2017) and splicing efficiency SE = 1-[mean intron coverage/mean coverage (last 30bp of upstream exon, first 30bp of downstream exon)] of nascent transcripts determined as described (Saldi et al. 2021).

### eNET-seq

eNET-seq was as described (Fong et al. 2022) with minor modifications. RNA decapping was with mouse decapping enzyme (NEB M0608S). Libraries were made using the Qiaseq miRNA Library kit (Qiagen 331505) with 12 base UMIs. Immunoprecipitation was with polyclonal rabbit anti pan-RNAPII CTD.

### eNET-seq read processing

Adapters were trimmed using cutadapt (v2.3) and reads were aligned to the hg38 human genome using Bowtie2 (v2.3.2). PCR duplicates were removed using UMI-tools (v0.5.4) and read coordinates were collapsed to a single base pair coordinate corresponding to the RNA 3’ end. Reads were filtered to only include those with a mapping quality score >=10 and to remove reads aligning to snoRNA genes. The relative signal (relative frequency, Fig. S1A) was calculated separately for each gene by dividing the signal in each bin by the sum of the signal for the entire plotted region. The relative signal was then averaged for each bin and multiplied by 1000 to remove small decimals. The mean relative signal was calculated separately for each biological replicate.

Pause sites were located by identifying positions where eNET-seq signal was >3 standard deviations above the mean for the surrounding 100 bp on each side, there were at least 5 reads at the pause site, and at least 5 additional reads within the window as previously described (Fong et al. 2022).

Pausing at 5’SSs was analyzed published eNETseq datasets (GSM6132196, GSM6132197, <u>GSM8313809</u>, <u>GSM8313810</u>, <u>GSM8313811</u> <u>GSM8313812</u>). Although Avitag Rpb1 has an Amr mutation, cells were not treated with α-amanitin.

## Data and code availability

RNA-seq, tNETseq, and eNET-seq data have been deposited at GEO.

## Acknowledgements

We thank S. Zhang MRC LMB, H. Shenasa (Stanford U.) and A. Fiszbein (Boston U) for valuable discussions and the UC Denver sequencing facility. D.B. thanks the Francis Crick Institute for their hospitality. Supported by NIH grant R35GM118051 to D.B., R35GM133385 to M.T. and R35GM145289 to R.Z.

## Author contributions

D.B. conceived the project with input from M.T., A.H., and R.X. N.F., R.S., M.T., A.H. and D.B. designed experiments. N.F. performed all experiments M.T, R.S. and N.F. performed bioinformatics. M.T. and D.B. wrote the paper with input from all authors.

